# A Comprehensive Annotation of Conserved Protein Domains in Human Endogenous Retroviruses

**DOI:** 10.1101/2025.07.25.666750

**Authors:** Tomàs Montserrat-Ayuso, Anna Esteve-Codina

## Abstract

Human endogenous retroviruses (HERVs) occupy nearly 8% of the human genome, yet their protein-coding potential remains largely unexplored. HERVs originate from ancestral exogenous retroviruses that infected germline cells and became integrated into the human genome. Like their exogenous counterparts, they typically follow the canonical proviral structure: LTR–gag–pol–env–LTR, where gag, pol, and env encode structural, enzymatic, and envelope proteins, respectively. Here, we present a comprehensive resource annotating conserved retroviral domains across 120,000+ ORFs derived from internal HERV regions. Using a reproducible pipeline based on HMMER and InterProScan, we identified over 17,000 domain hits—primarily from pol genes such as reverse transcriptase, RNase H, and protease—and quantified their structural conservation. Hundreds of domains exceed 95% alignment coverage, revealing a surprising abundance of full-length, retrovirus-like domains in both young and ancient HERV families. While the HERVK subfamily retains the most complete polyprotein architecture—including 13 loci with nearly intact Gag, Pol, and Env domains—many full-length Pol domains are also found in other families such as HERVH, HERVW, and HERVE. Our high-resolution annotations recover conserved catalytic motifs in Pol domains and transmembrane features in Env, enabling fine-grained functional interpretation. All annotations—including BED, FASTA, domain sequences, InterProScan outputs, and transmembrane predictions—are provided as an open resource for functional genomics and HERV expression studies at Zenodo (DOI: https://doi.org/10.5281/zenodo.17129661). This dataset will support downstream analyses of HERV protein expression, immune modulation, and co-option, in diseases and normal physiological conditions.

## Introduction

Endogenous retroviruses (ERVs) are remnants of ancient retroviral infections that became integrated into the germline and are now inherited as part of the host genome^1,2^. In humans, ERV (HERV) sequences account for approximately 8% of the genome and are typically found as fragmented long terminal repeats (LTRs), solo LTRs, and degraded internal regions^1,3,4^. Over the course of evolution, most HERVs have been progressively inactivated by accumulated mutations, deletions, and recombination events, resulting in a loss of their functional capacity^1,5,6^. Nonetheless, some HERV insertions have retained identifiable sequence features, such as intact open reading frames (ORFs) or conserved cis-regulatory motifs^6,7^. In several cases, these elements have been co-opted by the host genome to contribute to host physiological processes^8^. A well-known example is the syncytin gene family, derived from retroviral envelope (*env*) genes, which is essential for trophoblast fusion and placental morphogenesis in eutherian mammals^9,10^. These findings suggest that, despite widespread degradation, HERVs may still harbor biologically meaningful sequences, including those encoding functional protein domains.

The canonical retroviral genome encodes three major gene classes: *gag*, which produces capsid and matrix structural proteins; *pol*, encoding key enzymatic machinery including protease, reverse transcriptase, RNase H, and integrase; and *env*, which mediates viral entry through membrane fusion^1,11^. Although these genes are often fragmented or lost in endogenous retroviruses, several HERV loci retain partial or complete ORFs corresponding to these gene classes^5,12–14^. The biological relevance of these retained sequences is underscored by examples of co-option, such as the syncytins mentioned above, but also by growing evidence of HERV-derived proteins involved in immune modulation, neurodevelopment, and tumorigenesis^8,15–19^. Although most HERVs are considered to be usually dormant, they can be reactivated by stimuli such as viral infection, including influenza, HIV, or herpesviruses^15,20^, underscoring their dynamic interplay with the host immune system. This reinforces the importance of identifying conserved HERV ORFs and their associated retroviral domains as potential contributors to cellular processes or pathology.

While much recent research has focused on the transcriptional activity and regulatory roles of HERV-derived sequences—such as their enrichment in enhancer elements, influence on chromatin states, or contribution to long non-coding RNAs^7,21^—less attention has been paid to their protein-coding capacity, particularly at the domain level. Prior studies often focus on specific HERV families (e.g., HERVK or HERVW), or on single co-opted genes, rather than conducting genome-wide surveys of protein-coding domain conservation^8,9,22–26^. Moreover, to our knowledge no studies have systematically evaluated the structural integrity of identifiable retroviral domains using coverage metrics from hidden Markov model (HMM) alignments.

One close approximation is the analysis by Nakagawa and Takahashi^13^, which identified thousands of HERV-derived ORFs with partial or complete conservation of retroviral domains using their gEVE database^13^. While this pioneering effort provided a valuable early resource for exploring HERV coding potential, the gEVE pipeline was developed in 2016 and relies on software such as RetroTector, which can be difficult to integrate into modern reproducible workflows^27^. Currently, there is no community-standard tool for HERV discovery, and most available methods lack shareable and reproducible pipelines^27,28^. Furthermore, the gEVE framework did not quantify domain coverage or systematically examine subfamily-specific conservation patterns, limiting its ability to assess the structural integrity of retroviral genes at high resolution. Building upon these foundations, our study provides a reproducible, domain-centric analysis that enables finer-scale interrogation of protein-coding potential across the HERV landscape. Consequently, the extent to which HERVs retain well-structured protein domains across the human genome—particularly within the enzymatic regions of Pol—remains poorly understood. Furthermore, recent work has shown that even non-retroviral endogenous viral elements may retain protein domains with possible antiviral functions, underscoring the broader relevance of domain-level annotation in paleovirology and host-virus coevolution^29^.

Several resources have contributed to improving the classification, localization, and annotation of HERVs in the genome. The HERVd database was among the first systematic catalogs of HERV families and chromosomal coordinates, based on RepeatMasker and profile HMM-based annotation^30^. The above-mentioned gEVE database expanded this framework to multiple vertebrate genomes, integrating domain-level information with genomic context to support comparative analysis^13^. More recently, the Telescope pipeline introduced a high-resolution approach to locus-specific HERV expression quantification in RNA-seq data, including the development of a curated database of HERV loci and family-level annotations that has become foundational for expression-based studies^31^. Additionally, the ERVmap resource provided a locus-specific catalog of over 3,000 near full-length HERVs and a dedicated RNA-seq quantification pipeline, enabling high-resolution analysis of HERV expression across cell types and disease states, including autoimmune and cancer contexts^32^. These resources have significantly improved our understanding of HERV transcription and classification, but none have systematically addressed domain-level structural conservation in predicted ORFs.

In this study, we present a genome-wide analysis of the structural conservation of retroviral protein domains within ORFs derived from internal HERV sequences. Despite extensive mutational decay, we identify hundreds of HERV loci that retain high-confidence domains from the *gag*, *pol*, and *env* gene classes—many with complete or near-complete coverage of HMM profiles. While most studies examine HERVs at the subfamily level, this approach may obscure functional heterogeneity, as only a subset of loci within a given subfamily retain structurally intact domains. By quantifying domain-level conservation at single-locus resolution, we define a set of structurally preserved retroviral proteins that may represent candidates for residual biological activity or host co-option, providing a foundational resource for future functional investigations.

Our analysis builds upon previous resources such as gEVE and ERVmap^13,32^, shifting the focus from presence/absence annotation to domain-centric conservation with quantitative resolution. To our knowledge, this is the first genome-wide resource to systematically annotate ERV ORFs at the domain level, integrating alignment coverage, conserved motifs, and structural features into a shareable and reproducible dataset.

## Methods

### Repeat Masker annotation

We annotated human endogenous retroviral (HERV) insertions across the human genome using RepeatMasker^33^ version 4.1.8, employing NCBI/RMBLAST (v2.14.1+) as the search engine. The analysis was conducted on the GRCh38/hg38 reference genome (primary assembly) using the Dfam 3.9 repeat library^34^. RepeatMasker was run with -species human and -s and -no_is options.

### Extraction of Internal ERV Coordinates

To identify internal regions of human endogenous retroviruses (HERVs), we parsed RepeatMasker .out files for entries labeled with “-int” or “internal” in their name and classified as LTR/ERV. These entries were converted into 6-column BED format (chromosome, start, end, name, score, strand), where the name included chromosomal position, strand, and divergence from consensus.

### Merging of Overlapping Internal Regions

To reduce redundancy and better reflect transcript-like HERV structures, we merged overlapping or closely spaced internal HERV regions (≤150 bp gap) sharing the same subfamily, strand, and chromosome. Merging was performed grouping regions by subfamily and strand and collapsing them into unified intervals with updated standardized BED names.

### Sequence Extraction of Merged ERVs

We extracted strand-specific DNA sequences of the merged HERV internal regions using the pyfaidx Python package. Sequences were obtained from the indexed reference genome FASTA (GRCh38) and reverse-complemented if on the negative strand.

### ORF Prediction

ORFs were predicted from the merged internal HERV sequences using the EMBOSS getorf tool (v6.6.0)^35^. Sequences were provided in FASTA format, and ORFs shorter than 180 nucleotides were discarded using the -minsize 180 parameter. To maximize sensitivity, ORFs were defined from stop codon to stop codon, without requiring a canonical start codon, by using the -find 0 and -methionine N options. This approach, described in Villesen et al. (2004)^5^ accommodates the non-conventional translation mechanisms commonly used by retroviruses—such as ribosomal frameshifting and termination suppression—particularly at internal *pol* gene. Also, ORF sequences lacking a canonical ATG start codon may still serve as coding exons in spliced retroviral transcripts^13^. ORFs were extracted in the forward strand only (-reverse N), since strand orientation was previously accounted for during sequence preparation.

### Protein Domain Detection

Retroviral protein domains were detected in ORFs (translated to amino acid sequences) using hmmscan from the HMMER (v3.4) package^36^ against the profile database from GyDB^37^. Results were parsed and filtered to retain only hits satisfying the following criteria: full-sequence E-value ≤ 1e-5, domain E-value ≤ 1e-5. To avoid redundancy and focus on the most relevant domain instance per locus, we further filtered the results to retain only the best-scoring match per domain type (e.g., Gag, RT, RNaseH, Env) within each ORF. Domains were categorized into gene classes (e.g., *gag*, *pol*, *env*, *accessory*) according to GyDB classification. Accessory proteins are additional genes beyond the canonical *gag*, *pol*, and *env*. They are non-essential genes often involved in immune evasion, replication efficiency, or modulation of host functions^11^.

### Mapping Protein Domains Back to Genomic Coordinates

Filtered domain hits were mapped back to their corresponding genomic positions based on ORF-to-internal-sequence offsets and internal-to-genome coordinates. This conversion took into account strand orientation and translated amino acid coordinates into nucleotide positions. The mapping step produced BED-format annotations including the domain type, gene class, alignment score, and domain conservation level.

### Extraction of Domain Sequences

Protein domain coordinates from filtered HMMER output were used to extract precise amino acid subsequences corresponding to each domain. Domain sequences were retrieved from the original ORF FASTA based on alignment start and end positions, generating a FASTA file of domain-specific sequences for alignment and downstream analyses.

### Domain and motif annotation with InterProScan and Phobius

To complement HMMER-based domain detection, we annotated domain features and conserved motifs using InterProScan (v5.75)^38,39^. Protein domain sequences were analyzed using the InterProScan standalone tool with default parameters, enabling all member databases. The XML output was parsed to extract conserved sites predicted by the Conserved Domain Database^40^ (CDD), particularly focusing on catalytic residues and functional motifs. A custom Python script was used to extract and summarize the domain descriptions and conserved residues per protein.

To investigate membrane-associated features of Env proteins, we filtered the predicted ORFs to retain only sequences belonging to *env* genes. These sequences were then analyzed using Phobius^41^ with default settings to predict transmembrane domains and signal peptides. The output was parsed to identify Env proteins with predicted transmembrane regions, which may indicate preserved envelope protein topology.

The flow chart of the complete pipeline is shown in Figure 1 and it is available as a series of standalone scripts (see below and “Code availability”). All domain annotations, including BED files, FASTA sequences, and InterProScan outputs, were generated using this pipeline. These datasets are available at Zenodo (repository: “Domain-Level Annotations and Conservation Scores for Human Endogenous Retroviruses”, DOI: https://doi.org/10.5281/zenodo.17129661), and the Python scripts used to generate them are available for reproducibility at GitHub:https://github.com/funcgen/herv-domain-map.git.

**Figure 1.**
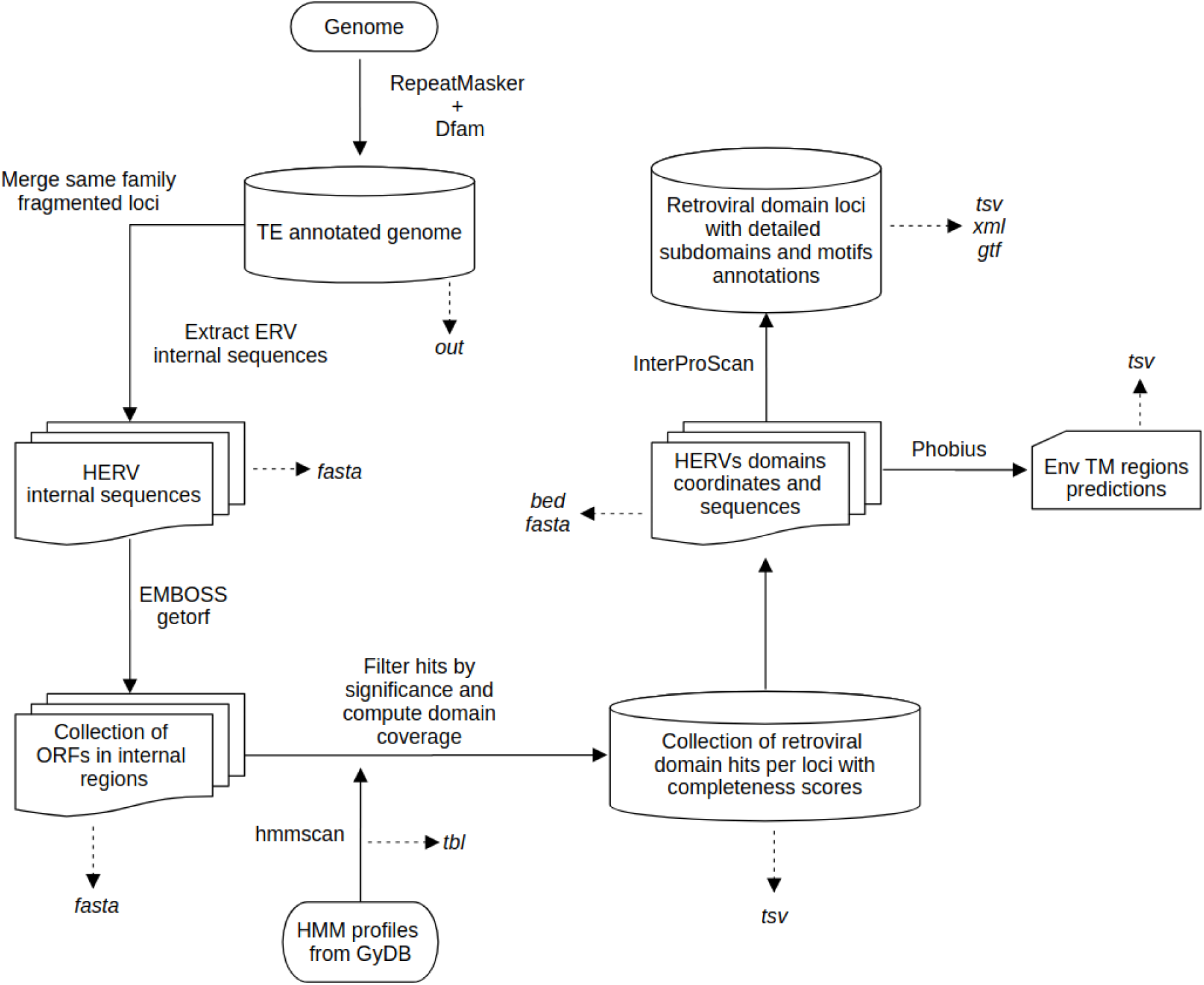
The annotation pipeline begins with the output file from RepeatMasker. While the main output is a TSV file containing domain coordinates and completeness scores, several intermediate and supporting files are also generated, including FASTA, TSV, BED, and TBL files, as well as InterProScan and Phobius output files providing additional information for each detected domain.

### Identification and Analysis of Conserved Protein Domains in HERV ORFs

Amino acid sequences were loaded in R using the Biostrings package and characterized by length and chromosomal distribution. Domains were classified as conserved if coverage was ≥40%, and as highly conserved if coverage exceeded 95%. ORFs were grouped by domain class, and the number of ORFs harboring conserved domains was summarized. Domain coverage distributions were visualized using violin and beeswarm plots, highlighting thresholds used for conservation categorization (40%, and 95%).

Subfamily-level analyses focused on four major HERV subfamilies (HERVK, HERVH, HERV17, and HERVE). Within each subfamily, the coverage scores of eight key domain types (Gag, RT, RNaseH, DUT, INT, AP, Env, and accessory) were assessed. For each ORF and domain class, only the best-scoring domain match was retained.

To evaluate co-occurrence of domain types within individual loci, we grouped ORFs by locus and determined the presence of Gag, Pol, and Env domains. Loci containing all three gene classes (i.e., “trios”) with conserved domains were flagged as potentially retaining multi-domain retroviral architecture. A stricter criterion requiring >80% coverage for each domain was also applied to define high-confidence candidate loci. For these loci, domain-specific coverage and subfamily information were compiled into summary tables for further inspection.

All statistical analyses and plots were performed in R (v4.3.3) using the dplyr^42^, ggplot2^43^, ggbeeswarm, Biostrings^44^, and openxlsx^45^ packages.

## Results

### High Prevalence of Conserved Domains in HERV ORFs

To explore the functional potential of endogenous retroviral sequences, we systematically annotated ORFs across the internal HERV sequences found by RepeatMasker^33^ and assessed the presence of conserved retroviral domains (see full annotation at Zenodo; repository: “Domain-Level Annotations and Conservation Scores for Human Endogenous Retroviruses”, DOI: https://doi.org/10.5281/zenodo.17129661). A total of 121,213 ORFs were identified and analyzed, with lengths ranging from 60 to 1415 nucleotides (median = 79 nt; mean = 92.89 nt). ORFs were distributed across all autosomes and sex chromosomes, with chr4 and chrX harboring the highest numbers (9,762 and 9,679 ORFs, respectively), while the fewest were observed on chr22 (934 ORFs). When normalized by chromosome size, distinct patterns emerged: the Y chromosome showed the highest ORF density, with one ORF every ∼10.4 kb, followed by chromosomes 19 and X. In contrast, chromosomes 16, 20, and 22 had the lowest densities (one ORF every 50–56 kb). As previously reported^3,46,47^, the accumulation of HERVs on the Y chromosome likely reflects reduced recombination, relaxed purifying selection, and a permissive environment for repetitive element retention. These observations point to a non-random genomic distribution of HERV-ORFs, potentially shaped by chromosome-specific differences in evolutionary constraint, integration bias, and retrotransposon dynamics.

Despite the short length of most ORFs, a total of 16,975 sequences still had identifiable retroviral-like domains, We defined conserved domains as those covering ≥40% of the corresponding HMM profile—a threshold selected to balance sensitivity and specificity, allowing detection of partially preserved domains while minimizing inclusion of highly degraded fragments likely to be biologically uninformative. Applying this criterion, we identified 6,501 domains with moderate to high conservation, each spanning hallmark gene classes including *gag*, *pol*, *env*, and accessory. The majority of hits corresponded to Pol-related domains (n = 5,891), which include key enzymatic components such as reverse transcriptase, integrase, RNase H, and aspartic protease. Fewer ORFs carried Env domains (n = 348), followed by Gag (n = 211), and accessory proteins (n = 51) (Fig. 2A). This distribution underscores the prevalence of conserved Pol-derived fragments in the genomic remnants of HERVs.

**Figure 2.**
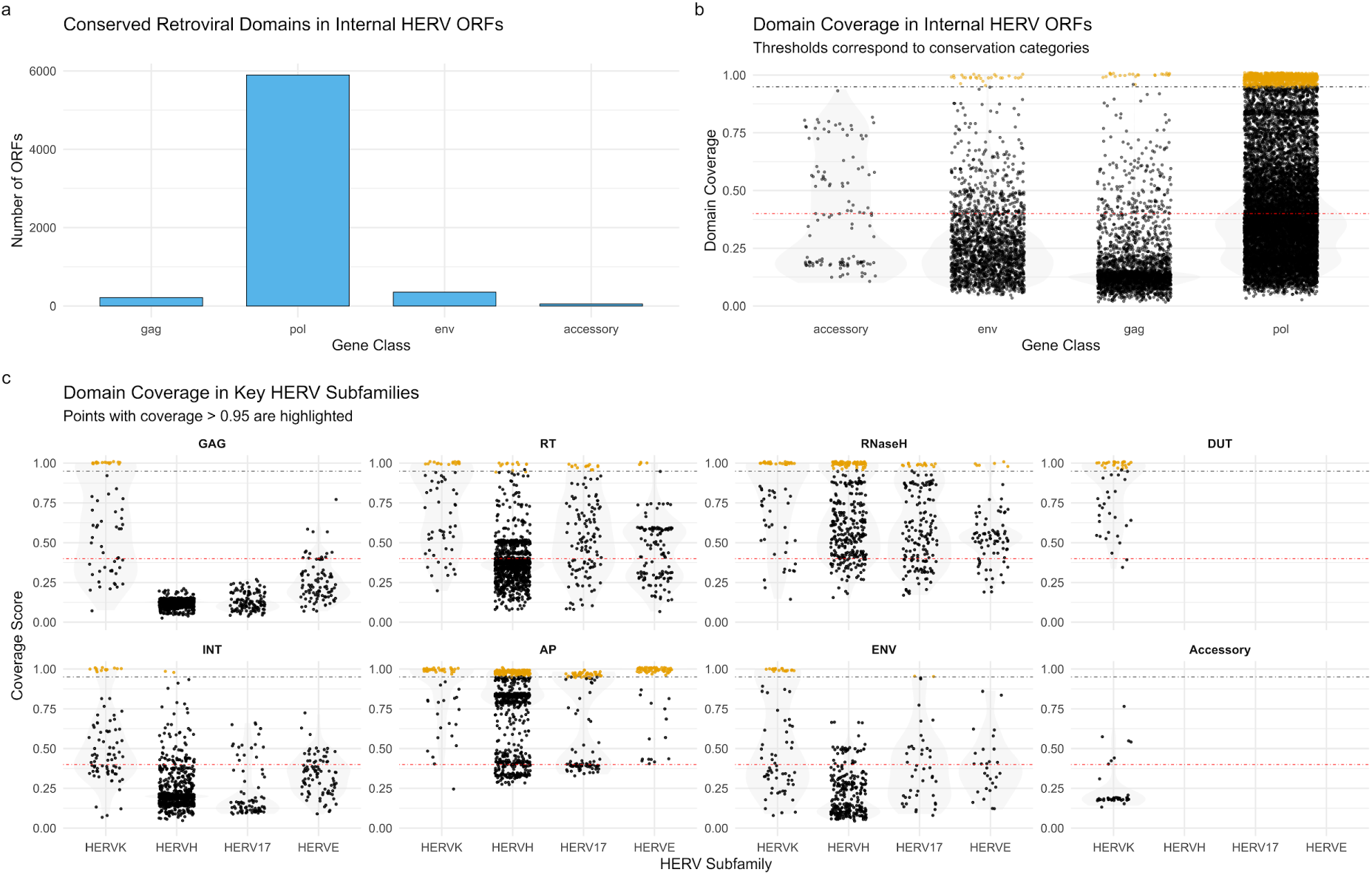
a) Barplot showing the number of annotated ORFs containing conserved domains (≥ 40 % HMM profile coverage) associated with each major retroviral gene class (*gag*, *pol*, *env*, and accessory). b) Violin plots showing the distribution of domain coverage values for annotated HERV ORFs grouped by gene class (accessory, env, gag, and pol). Each point represents an individual domain hit passing quality thresholds (e-value < 1e–5). The red dashed line marks the 40 % minimum domain coverage. Higher density of points near full coverage (1.0) is observed especially in the pol, gag and env classes, indicating a substantial number of highly conserved domains. Yellow points highlight full-length alignments (coverage > 0.95), reflecting potentially intact or structurally well-preserved retroviral sequences. c) Data points represent both total identified domains (black) and highly conserved domains (coverage > 0.95; highlighted). HERVK and HERVH exhibit the most extensive domain retention, while other subfamilies show gene-specific or restricted conservation.

The presence of these domains across a diverse set of chromosomal locations suggests widespread distribution and long-term retention of retroviral elements with partially conserved coding capacity.

These findings support the notion that while many HERVs are degenerated, a subset retains traces of protein-coding potential, particularly within the *pol* gene family.

### Substantial Fraction of Domains Show High Structural Conservation

To further characterize the quality of the annotated HERV-derived domains, we evaluated their structural conservation by analyzing alignment coverage against reference HMM profile. As mentioned above, a substantial proportion of domains exhibited high coverage values: many surpassed the 0.4 coverage threshold, with numerous domains reaching coverage values above 0.8 or even complete alignment (coverage = 1.0) (Fig 2B). These high-coverage matches suggest that several endogenous retroviral elements retain extensive homology to their exogenous retroviral counterparts.

Several representative examples illustrate the degree of conservation observed. One Env domain from a HERVK locus (chr5:156658763–156665917) showed 99.5% coverage over 356 amino acids, with an exceptionally high HMMER score (829.4) and a highly significant e-value (4.9e–251), strongly supporting its structural integrity. To further evaluate its functional conservation, we inspected the results from InterProScan and Phobius. The analysis revealed a complete GP41-like region, including conserved HR1–HR2 heptad repeats, a predicted transmembrane helix, and cytoplasmic and non-cytoplasmic domains (Supplementary Fig. 1). These structural features are essential for membrane fusion in retroviral envelope proteins, suggesting that this Env domain may retain key aspects of its ancestral function.

Another example comes from a Pol RNaseH domain encoded by a HERVH locus (chr14:53129175–53135122), which showed full coverage over 148 amino acids, with a high HMMER score (153.7) and a significant e-value (6.6e–47). InterProScan annotation identified a complete RNaseH_HI_RT_Bel domain, matching multiple conserved RNase H family signatures, including RNASE_H_1 (PFAM), RNaseH_domain (PROSITE), and RNaseH_sf (CATH-Gene3D). The region exhibited consistent hits across multiple databases, including PFAM, SUPERFAMILY, and CDD, supporting its classification as a structurally conserved RNase H-like fold. The conserved DEDD residues—characteristic of active RNase H enzymes^48^—were also detected, reinforcing the potential enzymatic activity of these domains. (Supplementary Fig. 2). Although catalytic activity was not directly tested, the presence of conserved residues typically associated with RNaseH function supports potential enzymatic functionality, possibly contributing to nucleic acid metabolism or retroelement regulation.

As a final example, we highlight a Pol protease domain from a HERV-E locus (chr1:20154322–20160102). Despite its shorter length (76 amino acids), this domain showed complete coverage of the HMM profile, with a strong HMMER score (67.5) and a significant e-value (1.4e–20). InterProScan identified a complete ASP_PROT_RETROV domain, supported by multiple overlapping annotations across PFAM (RVP – Retroviral Aspartyl Protease), PROSITE (Peptidase_A2_cat), SUPERFAMILY (Acid Proteases), and CATH-Gene3D (Peptidase_aspartic_dom_sf) (Supplementary Fig. 3). These consistent annotations indicate that the domain adopts the conserved structure of pepsin-like retroviral aspartyl proteases, suggesting structural retention in this ancient HERV lineage. Interestingly, the annotated gene ENSG00000227066 in Gencode^49^ (v48), described as an uncharacterized lncRNA, neatly overlaps this HERV, specifically the RNase H (full-length), and integrase (degraded) domains.

Beyond individual examples, we performed a systematic analysis of 1,006 full-length domains (HMMER coverage >95%), encompassing RNaseH, protease, integrase, reverse transcriptase, Env, Gag, and accessory proteins. Among these, 354 domain instances contained conserved catalytic or structural residues as annotated by InterProScan when searching in the CDD. RNaseH domains accounted for the majority of these cases, frequently retaining residues essential for enzymatic activity (e.g., conserved D, E, D, D) and RNA/DNA hybrid binding sites (161 instances retaining at least the latter, 5 retaining both). Similarly, 79 dUTPase and 72 protease domains showed well-conserved active or catalytic site residues. For Env domains, five full-length instances—belonging to the HERVS71, HERVW, and HERVE-a subfamilies—contained sequences that matched known immunosuppressive regions in the CDD. Complementary analysis with Phobius further revealed that full-length ENV domains frequently encode C-terminal transmembrane helices, reinforcing their structural completeness and potential for functionality.

These results highlight that, while many HERV loci are fragmented and degraded, a notable fraction retain high levels of sequence conservation in specific protein domains. This supports the potential relevance of these elements in host biology and evolutionary dynamics.

### Subfamily-Level Patterns of Domain Retention

To investigate the evolutionary preservation of endogenous retroviral proteins, we examined the distribution of conserved domains across HERV subfamilies (Supplementary Table 1 and Fig 2C). While some subfamilies showed widespread retention of the canonical retroviral genes *gag*, *pol*, and *env*, others exhibited more restricted or partial conservation.

The HERVK family displayed the most diverse and abundant repertoire of conserved domains, with multiple clades (e.g., HERVK, HERVK9, HERVK11, HERVK14, HERVK22) retaining domains from all major gene classes: *gag*, *pol*, *env*, and in some cases, accessory genes. For example, HERVK9 alone contributed over 100 hits each for aspartic protease (AP), reverse transcriptase (RT), and RNase H domains. Even under stringent filtering (coverage > 0.95, Supplementary Table 2), HERVK retained a notable number of full-length domains — including 33 AP, 28 RNaseH, 27 dUTPase (DUT), 17 RT, 13 integrase, 17 Gag, and 19 Env — indicating preservation of near-complete polyproteins in several loci.

In contrast, HERVH showed a clear Pol-centric conservation pattern. It contributed the highest number of AP (n = 700), RNase H (n = 308), and RT (n = 307) domains across all subfamilies (≥ 40 % coverage), but lacked any conserved GAG domains and retained only 35 ENV hits. Under the high-confidence threshold, it preserved 185 full-length AP, 76 RNase H, 13 RT and 2 integrase domains, highlighting a selective retention of enzymatic functions and suggesting pressure against structural gene conservation.

The HERVW lineage, primarily represented by HERV17^3^, exhibited moderate but consistent conservation of Pol, and Env domains. It retained dozens of hits across Pol domains, and under the >0.95 coverage threshold, 51 AP, 17 RNase H, 14 RT, and 2 ENV domains remained — supporting a degree of functional maintenance within this subfamily.

Several other lineages showed more specialized conservation patterns. HERVE and HERV9, for instance, contributed primarily Pol-derived domains (especially proteases), while Env-enriched conservation was mostly restricted to lineages like HERVIP10FH or Harlequin. Notably, HERVE — considered evolutionarily ancient — retained 69 full-length AP domains, 6 RNase H, and 4 RT domains, suggesting unexpected structural integrity.

Overall, the conservation of full-length domains in specific subfamilies, including both evolutionarily young (e.g., HERVK) and ancient (e.g., HERVE) lineages, highlights the selective retention of replication-related proteins. This pattern of domain conservation raises the possibility that some HERVs may preserve structural or functional potential long after their integration.

### Co-occurrence of Conserved Retroviral Domains Reveals Residual Proviral Architecture

The presence of all three domain types—Gag, Pol, and Env—is essential for the formation of retroviral particles, as they respectively encode structural components, enzymatic machinery for reverse transcription and integration, and the envelope proteins required for cell entry.

To assess the potential for residual functionality among HERV loci, we examined the co-occurrence of conserved domains within each HERV insertion. We first identified loci that contain at least one Gag, one Pol, and one Env domain, without applying a stringent conservation threshold (only 40% of domain coverage). Such configurations are a prerequisite for viral-like particle formation. Under this relaxed criterion, 41 loci were found to carry a combination of structural and enzymatic domains, suggesting the preservation of core retroviral architecture at a broad scale (Supplementary Table 3).

To prioritize high-confidence candidates, we applied a stricter filter requiring all three domain types (Gag, Pol, and Env) to have at least one domain of each with a minimum alignment coverage of 0.8. This refinement yielded 13 loci, all belonging to the HERVK subfamily, that retained a full complement of core retroviral domains with high sequence conservation (Table 1 and Supplementary Table 4).

**Table 1.**
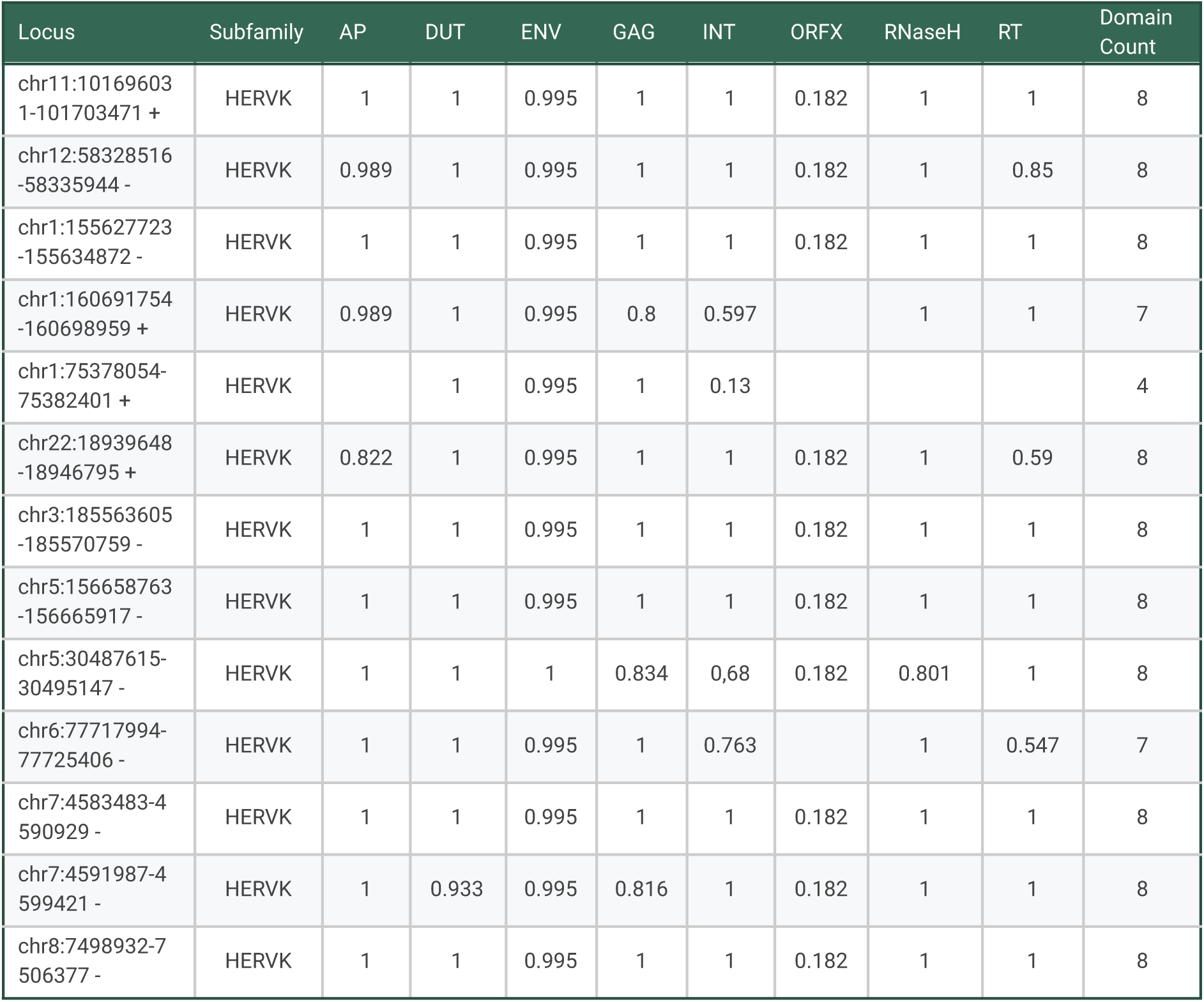
Top HERVK loci with co-occurring Gag, Pol, and Env domains (coverage > 0.8 in at least one domain). Each locus encodes seven distinct retroviral domains with high alignment coverage, except one locus, encoding 3.

These 13 loci typically encode up to seven distinct domains, encompassing aspartic protease (AP), dUTPase (DUT), reverse transcriptase (RT), RNase H, integrase (INT), Env, and Gag. Most domains exhibited near-complete alignment (coverage > 0.95), with several loci achieving full-length coverage across all seven domains. All 13 loci retained both 5′ and 3′ LTRs, consistent with full-length proviral structure. For example, the insertion at chr1:155627723–155634872 shows perfect conservation (coverage = 0.995-1.0) for every domain. Similar complete or near-complete configurations were observed on chromosomes 3, 5, 7, 8, 11, and 12 (Table 1 and Supplementary Table 3). Interestingly enough, several of these HERVK loci are located within introns of human genes (e.g., *SGCD*, *MEI4*, *DEFB107B*), often in the antisense orientation, suggesting possible regulatory interactions.

These loci represent the best candidates for retained protein-coding potential among endogenous retroviruses and capacity to form infectious viral-like particles. Their intact domain architecture may reflect recent integration, selective constraint, or co-option into host regulatory or structural functions. The functional and structural potential of these elements motivates further studies into their transcriptional activity, epigenetic regulation, and potential immunogenicity, and, notable, two loci (e.g., chr1:155627723–155634872) encode Env in the same ORF as several Pol domains, an unusual configuration that may reflect evolutionary fusion events.

## Discussion

Our systematic analysis reveals that thousands of ORFs embedded within HERV loci encode recognizable retroviral protein domains, despite the extensive genomic erosion these elements typically undergo. Many of these ORFs exhibit high domain coverage and strong alignment scores, indicating the retention of structural features beyond what would be expected from random degradation. Notably, we observe unexpectedly high conservation within Pol-derived domains, including reverse transcriptase, RNase H, integrase, and aspartic protease. Several RNase H and protease domains align fully or nearly fully to their respective HMM profiles and retain catalytic motifs such as the DEDD residues in RNase H domains, a hallmark of enzymatic activity^48^. While prior studies have described isolated ORFs or co-opted genes for *env* or *gag*^5,8,50^, our genome-wide approach extends this view by systematically quantifying domain-level conservation across all major HERV subfamilies, including Pol domains. These findings suggest that HERV-derived *pol* genes may represent a broader and more structurally intact source of retroviral legacy than previously recognized.

The conservation of protein domains is not uniformly distributed across HERV lineages, but instead reflects subfamily-specific patterns. Among these, the youngest HERVK family, as expected^3^, retains multiple loci with near-complete retroviral architecture—including co-occurring Gag, Pol, and Env domains—in configurations reminiscent of intact retroviruses. This is consistent with reports of active HERVK expression and translation in early embryonic tissues and cancer^21,51–53^. In contrast, HERVH rarely retains intact Gag and Env ORFs—*env* in particular—, yet exhibits striking conservation of Pol-encoded enzymatic domains, especially RNase H and protease. This pattern aligns with previous findings indicating extensive recombination and truncation in HERVH loci, which have been maintained at high copy number due to regulatory or transcriptomic utility^7,54^. Our results refine this view by showing that, despite structural erosion, a subset of HERVH elements preserves enzymatic domain integrity—suggesting selective maintenance of Pol-related functions (specially RNase H and aspartic protease activity). These contrasting patterns underscore how domain retention may reflect differential evolutionary pressures, with some domains retained for potential functional co-option, even in the absence of full retroviral structure, possibly enabling domain-level co-option or, in some contexts, trans-complementation.

Although most HERV loci are fragmented and non-functional due to accumulated mutations, our analysis reveals that retroviral protein domains exhibit variable levels of structural conservation across loci. In many cases, no single locus retains all three canonical domains in full. However, the presence of complementary domain preservation at distinct genomic sites raises the possibility of functional trans-complementation, whereby transcripts or proteins encoded by separate HERV loci could, in theory, assemble into a partially functional retroviral-like complex. This modular conservation suggests a distributed potential for retroelement activity, particularly if such loci are co-expressed in the same cellular context. While trans-complementation has been proposed as a mechanism to increase retroelement copy number^3^, it is generally considered rare in the context of HERVs^55^. Nevertheless, under conditions of widespread HERV reactivation—such as those observed in certain diseases^21^—this mechanism cannot be ruled out as a possible contributor to the formation of viral-like particles.

Beyond the potential for trans-complementation, the domain-level preservation also opens the door to alternative functional scenarios. In particular, the widespread conservation of enzymatic domains—especially RNase H and aspartic protease—in ancient subfamilies^6^ such as HERVH and HERVE raises the possibility of residual activity or structural co-option. Given the established role of retroviral proteases in polyprotein maturation^56^, their persistence may reflect functional retention or repurposing in host cellular contexts. Such conservation of retroviral enzymatic functions echoes known cases of co-option, such as the syncytins, which not only mediate placental fusion but also possess immunosuppressive properties^9,57^. Similarly, the preservation of full-length Env domains in several HERVK loci supports a potential antiviral role through receptor interference, a mechanism initially proposed in koalas and bats whereby Env blocks viral entry by binding host receptors^15,48^. In this context, it is conceivable that Pol-derived enzymatic domains such as RNase H and aspartic protease may also contribute to antiviral defense through dominant-negative interference or residual catalytic activity. For instance, HERV-derived RNase H might degrade viral RNA–DNA hybrids during reverse transcription, or modulate immune sensing by processing nucleic acid hybrids that accumulate in the cytoplasm. In a similar manner, residual protease activity could interfere with the processing of viral polyproteins. These actions would mirror established antiviral strategies such as receptor interference by HERV Env proteins, and would conceptually parallel restriction factors such as TRIM5α and tetherin^58,59^, which block replication by binding and disrupting key viral processes.

Although direct experimental validation remains lacking, the conservation of these catalytic folds across ancient HERV lineages supports the notion that HERV proteins may persist not merely as evolutionary remnants, but as active participants in host-virus interactions.

Beyond potential residual function, our findings also support a broader evolutionary perspective: that HERVs, long dismissed as genomic “junk,” may have been retained in part due to their structural and regulatory utility^15,60–64^. Rather than being purely parasitic, these sequences can act as white sheets of genomic material—regions with the capacity to incorporate, rearrange, and express novel DNA. In this view, HERVs offer evolutionary scaffolds from which new genes, transcripts, or regulatory networks may emerge. The retention of coding domains in even ancient subfamilies suggests that this material may not only be tolerated by the genome, but in some cases, positively selected as a source of innovation.

Nevertheless, several important limitations must be considered. Our analysis relies solely on computational predictions of protein domains within candidate ORFs and lacks direct experimental evidence of expression or activity. We did not assess whether the identified ORFs are transcribed, translated, or post-translationally modified in vivo. Additionally, no functional assays were conducted to validate enzymatic activity or fusogenic potential. Therefore, while the structural conservation we observe provides strong evidence for potential functionality, empirical validation—including transcriptomics, proteomics, and targeted assays—is essential to fully assess biological relevance.

To address these open questions, future work should integrate domain annotations with bulk and single-cell transcriptomic data to identify actively expressed HERV loci. Ribosome profiling and mass spectrometry can further clarify which ORFs are translated and potentially functional. Moreover, structure prediction tools such as AlphaFold^65^ may help assess whether conserved domains adopt native-like folds compatible with enzymatic or structural roles. Comparative analyses across primates and other mammals could reveal lineage-specific patterns of domain retention and identify cases of convergent co-option. Finally, functional studies should explore whether these HERV-derived domains participate in host immunity, neurodevelopment, placentation, or other biological pathways where HERVs are increasingly implicated^15–18,48^. HERVs can modulate innate immunity by activating cytosolic sensors (e.g., RIG-I, MDA5) or TLR pathways such as TLR3, TLR9, and TLR4, particularly via Env proteins like HERV-W ENV and by RNA-DNA hybrids^15^. Env proteins have been shown to induce inflammatory cytokines including IL-1β and TNF-α, and may contribute to antiviral defense or pathology, as proposed in multiple sclerosis^15,66^. Together, these efforts will help determine whether the sequences we have identified are evolutionary relics—or underrecognized components of human molecular biology.

In conclusion, our study provides the first genome-wide map of structurally conserved HERV protein domains in humans and identifies hundreds of loci with potential for residual function or evolutionary co-option. These findings suggest that HERV-derived sequences may retain structural integrity that enables functional activity, with potential implications for both physiology and pathophysiology. By pinpointing specific candidates that preserve retroviral enzymatic or structural architecture, we offer a foundation for future functional and evolutionary studies into the enduring impact of endogenous retroviruses on human biology.

## Data availability

The human genome assembly GRCh38 can be downloaded from the GENCODE website: https://www.gencodegenes.org/human/. The HMM profiles used for domain annotation are available from the GyDB database: https://gydb.org/index.php?title=Collection_HMM. The full HERV domain annotation dataset, including BED and FASTA files, InterProScan outputs, and supporting scripts, is available via Zenodo at DOI: https://doi.org/10.5281/zenodo.17129661.

## Code availability

The pipeline used to identify open reading frames, annotate retroviral domains and analyze the results is available as a collection of Python, Bash, and R scripts at https://github.com/funcgen/herv-domain-map.git.

## Author contributions

T.M-A and A.E-C conceived the project. T.M-A performed the bioinformatic analyses. T.M-A and A.E-C wrote the manuscript. A.E-C supervised the project.

## Acknowledgements

We would like to thank our colleague Jessica Gómez-Garrido, from the Genome Assembly and Annotation team at CNAG, for her valuable advice during the development of the annotation pipeline.

## References

1. Griffiths, D. J. Endogenous retroviruses in the human genome sequence. Genome Biol. 2, REVIEWS1017 (2001).

2. Stoye, J. P. Endogenous retroviruses: Still active after all these years? Curr. Biol. 11, R914–R916 (2001).

3. Bannert, N. & Kurth, R. The evolutionary dynamics of human endogenous retroviral families. Annu. Rev. Genomics Hum. Genet. 7, 149–173 (2006).

4. Lander, E. S. et al. Initial sequencing and analysis of the human genome. Nature 409, 860–921 (2001).

5. Villesen, P., Aagaard, L., Wiuf, C. & Pedersen, F. S. Identification of endogenous retroviral reading frames in the human genome. Retrovirology 1, 32 (2004).

6. Ueda, M. T. et al. Comprehensive genomic analysis reveals dynamic evolution of endogenous retroviruses that code for retroviral-like protein domains. Mob. DNA 11, 29 (2020).

7. Ito, J. et al. Systematic identification and characterization of regulatory elements derived from human endogenous retroviruses. PLOS Genet. 13, e1006883 (2017).

8. Wang, J. & Han, G.-Z. Frequent Retroviral Gene Co-option during the Evolution of Vertebrates. Mol. Biol. Evol. 37, 3232–3242 (2020).

9. Mi, S. et al. Syncytin is a captive retroviral envelope protein involved in human placental morphogenesis. Nature 403, 785–789 (2000).

10. Cornelis, G. et al. Retroviral envelope gene captures and syncytin exaptation for placentation in marsupials. Proc. Natl. Acad. Sci. U. S. A. 112, E487–496 (2015).

11. Retroviruses. (Cold Spring Harbor Laboratory Press, Cold Spring Harbor (NY), 1997).

12. de Parseval, N., Lazar, V., Casella, J.-F., Benit, L. & Heidmann, T. Survey of Human Genes of Retroviral Origin: Identification and Transcriptome of the Genes with Coding Capacity for Complete Envelope Proteins. J. Virol. 77, 10414–10422 (2003).

13. Nakagawa, S. & Takahashi, M. U. gEVE: a genome-based endogenous viral element database provides comprehensive viral protein-coding sequences in mammalian genomes. Database J. Biol. Databases Curation 2016, baw087 (2016).

14. Vargiu, L. et al. Classification and characterization of human endogenous retroviruses; mosaic forms are common. Retrovirology 13, 7 (2016).

15. Srinivasachar Badarinarayan, S. & Sauter, D. Switching Sides: How Endogenous Retroviruses Protect Us from Viral Infections. J. Virol. 95, e02299–20 (2021).

16. Duarte, R. R. R., Nixon, D. F. & Powell, T. R. Ancient viral DNA in the human genome linked to neurodegenerative diseases. Brain. Behav. Immun. 123, 765–770 (2025).

17. Duarte, R. R. R. et al. Integrating human endogenous retroviruses into transcriptome-wide association studies highlights novel risk factors for major psychiatric conditions. Nat. Commun. 15, 3803 (2024).

18. Küry, P. et al. Human Endogenous Retroviruses in Neurological Diseases. Trends Mol. Med. 24, 379–394 (2018).

19. Stricker, E., Peckham-Gregory, E. C. & Scheurer, M. E. HERVs and Cancer-A Comprehensive Review of the Relationship of Human Endogenous Retroviruses and Human Cancers. Biomedicines 11, 936 (2023).

20. Evans, E. F., Saraph, A. & Tokuyama, M. Transactivation of Human Endogenous Retroviruses by Viruses. Viruses 16, 1649 (2024).

21. Wang, J., Lu, X., Zhang, W. & Liu, G.-H. Endogenous retroviruses in development and health. Trends Microbiol. 32, 342–354 (2024).

22. Ono, R. et al. Deletion of Peg10, an imprinted gene acquired from a retrotransposon, causes early embryonic lethality. Nat. Genet. 38, 101–106 (2006).

23. Matsui, T. et al. SASPase regulates stratum corneum hydration through profilaggrin-to-filaggrin processing. EMBO Mol. Med. 3, 320–333 (2011).

24. Nakaya, Y., Koshi, K., Nakagawa, S., Hashizume, K. & Miyazawa, T. Fematrin-1 Is Involved in Fetomaternal Cell-to-Cell Fusion in Bovinae Placenta and Has Contributed to Diversity of Ruminant Placentation. J. Virol. 87, 10563–10572 (2013).

25. Pastuzyn, E. D. et al. The Neuronal Gene Arc Encodes a Repurposed Retrotransposon Gag Protein that Mediates Intercellular RNA Transfer. Cell 172, 275–288.e18 (2018).

26. Ashley, J. et al. Retrovirus-like Gag Protein Arc1 Binds RNA and Traffics across Synaptic Boutons. Cell 172, 262–274.e11 (2018).

27. Ritsch, M., Brait, N., Harvey, E., Marz, M. & Lequime, S. Endogenous viral elements: insights into data availability and accessibility. Virus Evol. 10, veae099 (2024).

28. Goubert, C. et al. A beginner’s guide to manual curation of transposable elements. Mob. DNA 13, 7 (2022).

29. Belyi, V. A., Levine, A. J. & Skalka, A. M. Unexpected Inheritance: Multiple Integrations of Ancient Bornavirus and Ebolavirus/Marburgvirus Sequences in Vertebrate Genomes. PLOS Pathog. 6, e1001030 (2010).

30. Pačes, J., Pavlíček, A. & Pačes, V. HERVd: database of human endogenous retroviruses. Nucleic Acids Res. 30, 205–206 (2002).

31. Bendall, M. L. et al. Telescope: Characterization of the retrotranscriptome by accurate estimation of transposable element expression. PLOS Comput. Biol. 15, e1006453 (2019).

32. Tokuyama, M. et al. ERVmap analysis reveals genome-wide transcription of human endogenous retroviruses. Proc. Natl. Acad. Sci. 115, 12565–12572 (2018).

33. Tarailo-Graovac, M. & Chen, N. Using RepeatMasker to identify repetitive elements in genomic sequences. Curr. Protoc. Bioinforma. Chapter 4, 4.10.1–4.10.14 (2009).

34. Storer, J., Hubley, R., Rosen, J., Wheeler, T. J. & Smit, A. F. The Dfam community resource of transposable element families, sequence models, and genome annotations. Mob. DNA 12, 2 (2021).

35. Rice, P., Longden, I. & Bleasby, A. EMBOSS: the European Molecular Biology Open Software Suite. Trends Genet. TIG 16, 276–277 (2000).

36. Potter, S. C. et al. HMMER web server: 2018 update. Nucleic Acids Res. 46, W200–W204 (2018).

37. Llorens, C. et al. The Gypsy Database (GyDB) of mobile genetic elements: release 2.0. Nucleic Acids Res. 39, D70–74 (2011).

38. Blum, M. et al. The InterPro protein families and domains database: 20 years on. Nucleic Acids Res. 49, D344–D354 (2021).

39. Jones, P. et al. InterProScan 5: genome-scale protein function classification. Bioinformatics 30, 1236–1240 (2014).

40. Wang, J. et al. The conserved domain database in 2023. Nucleic Acids Res. 51, D384–D388 (2023).

41. Käll, L., Krogh, A. & Sonnhammer, E. L. L. A combined transmembrane topology and signal peptide prediction method. J. Mol. Biol. 338, 1027–1036 (2004).

42. Wickham, H., François, R., Henry, L., Müller, K. & Vaughan, D. Dplyr: A Grammar of Data Manipulation. (2025).

43. Wickham, H. Ggplot2: Elegant Graphics for Data Analysis. (Springer-Verlag New York, 2016).

44. Pagès, H., Aboyoun, P., Gentleman, R. & DebRoy, S. Biostrings: Efficient Manipulation of Biological Strings. (2025).

45. Schauberger, P. & Walker, A. Openxlsx: Read, Write and Edit Xlsx Files. (2025).

46. Flockerzi, A., Burkhardt, S., Schempp, W., Meese, E. & Mayer, J. Human Endogenous Retrovirus HERV-K14 Families: Status, Variants, Evolution, and Mobilization of Other Cellular Sequences. J. Virol. 79, 2941–2949 (2005).

47. Skaletsky, H. et al. The male-specific region of the human Y chromosome is a mosaic of discrete sequence classes. Nature 423, 825–837 (2003).

48. Moelling, K., Broecker, F., Russo, G. & Sunagawa, S. RNase H As Gene Modifier, Driver of Evolution and Antiviral Defense. Front. Microbiol. 8, (2017).

49. Mudge, J. M. et al. GENCODE 2025: reference gene annotation for human and mouse. Nucleic Acids Res. 53, D966–D975 (2025).

50. de Parseval, N., Lazar, V., Casella, J.-F., Benit, L. & Heidmann, T. Survey of human genes of retroviral origin: identification and transcriptome of the genes with coding capacity for complete envelope proteins. J. Virol. 77, 10414–10422 (2003).

51. Li, Z. et al. Expression of HERV-K correlates with status of MEK-ERK and p16INK4A-CDK4 pathways in melanoma cells. Cancer Invest. 28, 1031–1037 (2010).

52. Zhao, J. et al. Expression of Human Endogenous Retrovirus Type K Envelope Protein is a Novel Candidate Prognostic Marker for Human Breast Cancer. Genes Cancer 2, 914–922 (2011).

53. Rycaj, K. et al. Cytotoxicity of human endogenous retrovirus K-specific T cells toward autologous ovarian cancer cells. Clin. Cancer Res. Off. J. Am. Assoc. Cancer Res. 21, 471–483 (2015).

54. Jern, P., Sperber, G. O. & Blomberg, J. Definition and variation of human endogenous retrovirus H. Virology 327, 93–110 (2004).

55. Magiorkinis, G., Gifford, R. J., Katzourakis, A., De Ranter, J. & Belshaw, R. Env-less endogenous retroviruses are genomic superspreaders. Proc. Natl. Acad. Sci. 109, 7385–7390 (2012).

56. Spall, V. E., Shanks, M. & Lomonossoff, G. P. Polyprotein Processing as a Strategy for Gene Expression in RNA Viruses. Semin. Virol. 8, 15–23 (1997).

57. Muir, A., Lever, A. & Moffett, A. Expression and Functions of Human Endogenous Retroviruses in the Placenta: An Update. Placenta 25, S16–S25 (2004).

58. Stremlau, M. et al. The cytoplasmic body component TRIM5α restricts HIV-1 infection in Old World monkeys. Nature 427, 848–853 (2004).

59. Neil, S. J. D., Zang, T. & Bieniasz, P. D. Tetherin inhibits retrovirus release and is antagonized by HIV-1 Vpu. Nature 451, 425–430 (2008).

60. Smit, A. F. Interspersed repeats and other mementos of transposable elements in mammalian genomes. Curr. Opin. Genet. Dev. 9, 657–663 (1999).

61. Garcia-Perez, J. L., Widmann, T. J. & Adams, I. R. The impact of transposable elements on mammalian development. Dev. Camb. Engl. 143, 4101–4114 (2016).

62. Platt, R. N., Vandewege, M. W. & Ray, D. A. Mammalian transposable elements and their impacts on genome evolution. Chromosome Res. Int. J. Mol. Supramol. Evol. Asp. Chromosome Biol. 26, 25–43 (2018).

63. Nishihara, H. Transposable elements as genetic accelerators of evolution: contribution to genome size, gene regulatory network rewiring and morphological innovation. Genes Genet. Syst. 94, 269–281 (2020).

64. Johnson, W. E. Origins and evolutionary consequences of ancient endogenous retroviruses. Nat. Rev. Microbiol. 17, 355–370 (2019).

65. Jumper, J. et al. Highly accurate protein structure prediction with AlphaFold. Nature 596, 583–589 (2021).

66. Christensen, T. Association of human endogenous retroviruses with multiple sclerosis and possible interactions with herpes viruses. Rev. Med. Virol. 15, 179–211 (2005).

