## Supplemental Figures for "A Comprehensive Annotation of Conserved Protein Domains in Human Endogenous Retroviruses"

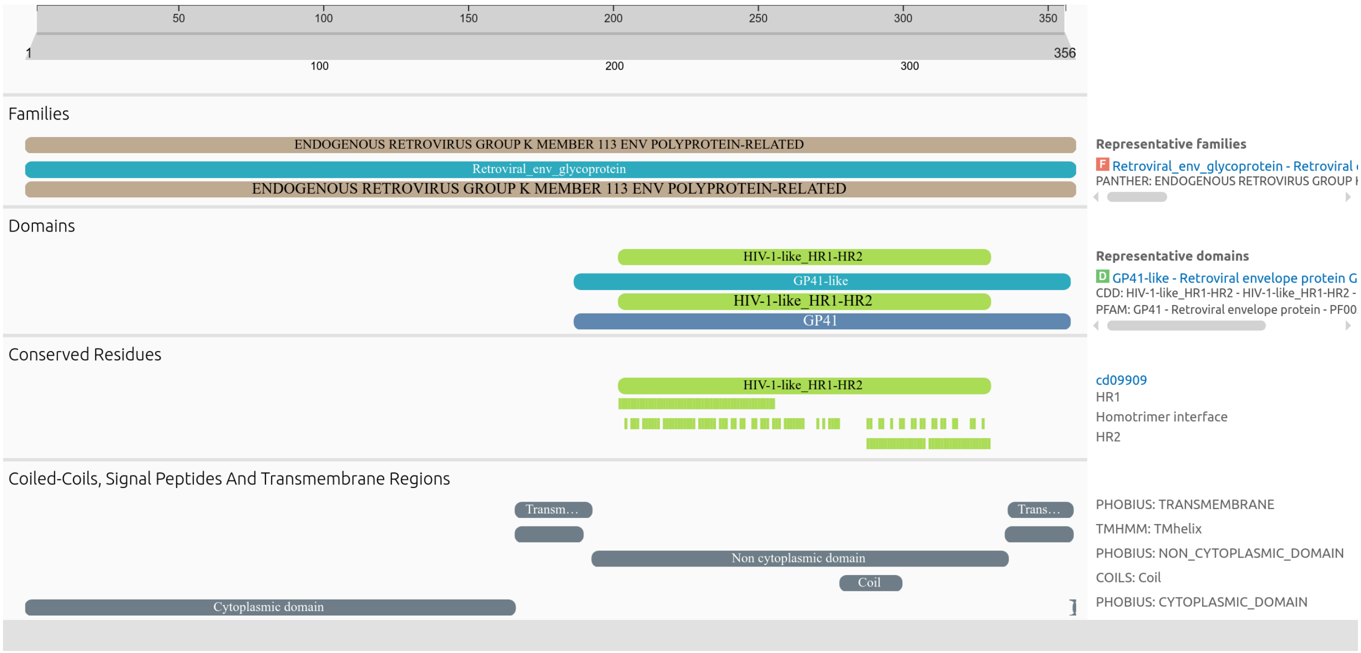


**Supplementary Figure 1.** *InterProScan annotation of a full-length HERV-K ENV domain (356 amino acids).*

The protein shows strong similarity to the retroviral envelope glycoprotein GP41 family, including conserved domains (HR1-HR2), transmembrane regions, and cytoplasmic/non-cytoplasmic boundaries. Green bars mark conserved residues. These features suggest retention of structural elements necessary for membrane fusion and Env function.

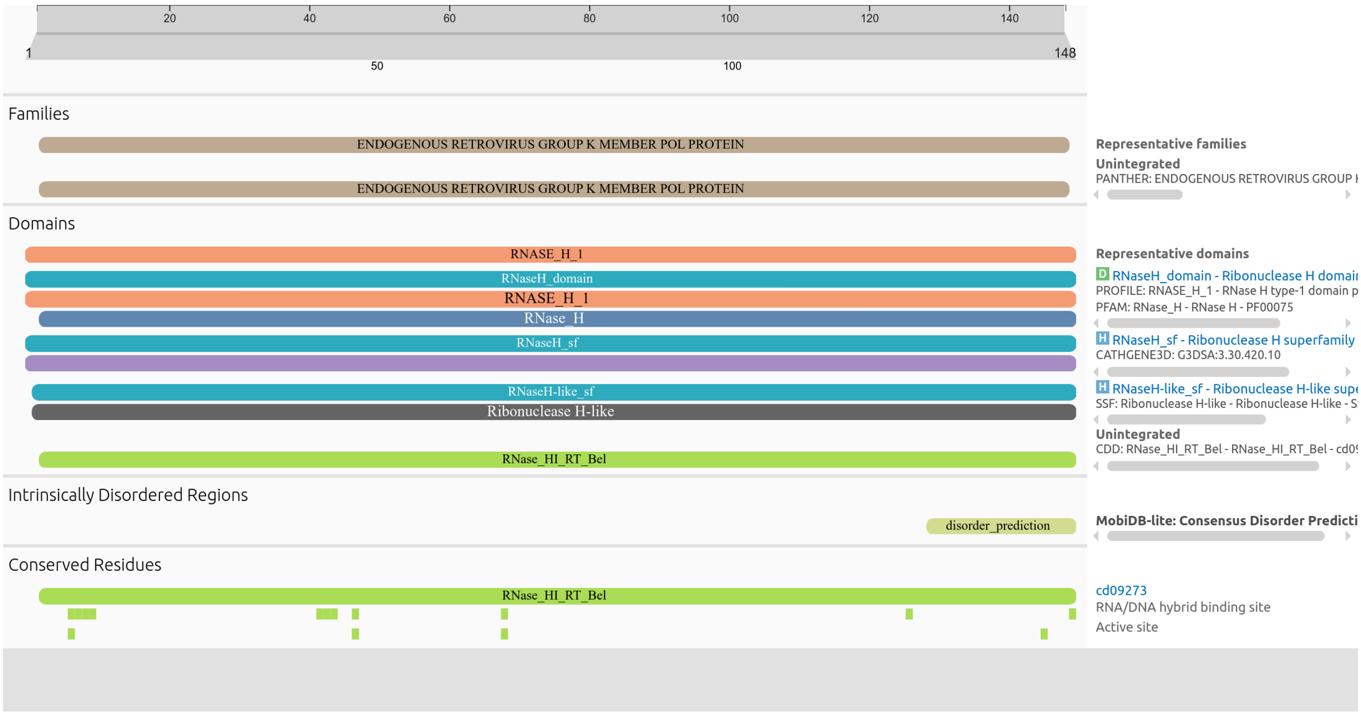


**Supplementary Figure 2.** *InterProScan annotation of a full-length HERV-H RNase H domain (142 amino acids).*

The domain is recognized by multiple families and databases, including PFAM, CDD, and CATH, and shows strong sequence conservation. Green bars highlight conserved residues, including active site positions and RNA/DNA hybrid binding motifs. This suggests potential retention of enzymatic function in the RNase H domain of this HERV element.

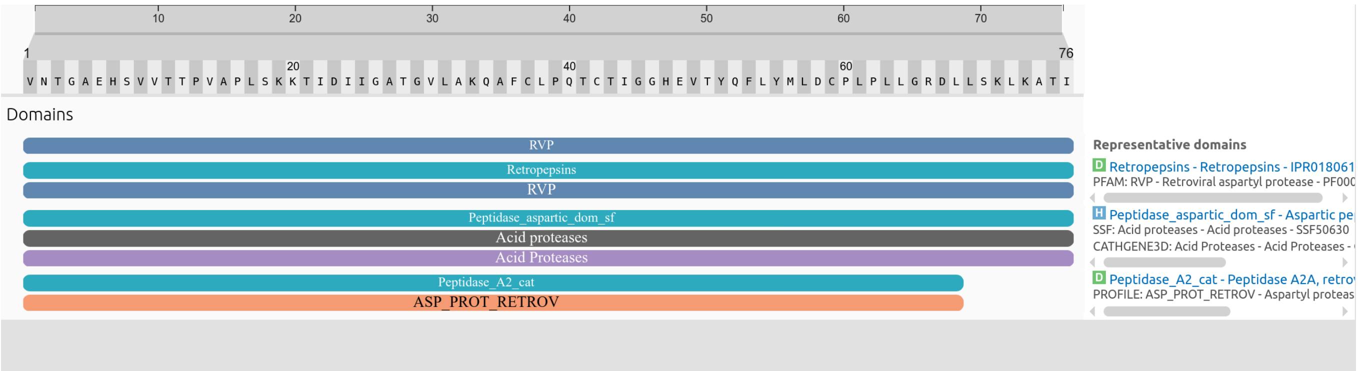


**Supplementary Figure 3.** *InterProScan annotation of a conserved HERV-E protease domain (76 amino acids).*

The protein displays multiple overlapping annotations corresponding to retroviral aspartyl proteases, including RVP (PFAM), ASP_PROT_RETROV (PROSITE), and Peptidase_A2_cat. Additional hits to acid protease superfamilies (CATH-Gene3D, SUPERFAMILY) further support structural conservation. These features indicate preservation of the characteristic fold of retroviral proteases, suggesting potential retention of ancestral proteolytic architecture in this endogenous retroviral locus.
